# Female behavior drives the formation of distinct social structures in C57BL/6J versus wild-derived outbred mice in field enclosures

**DOI:** 10.1101/2022.04.19.488643

**Authors:** Caleb C. Vogt, Matthew N. Zipple, Daniel D. Sprockett, Caitlin H. Miller, Summer X. Hardy, Matthew K. Arthur, Adam M. Greenstein, Melanie S. Colvin, Lucie M. Michel, Andrew H. Moeller, Michael J. Sheehan

**Affiliations:** Laboratory for Animal Social Evolution and Recognition, Department of Neurobiology and Behavior, Cornell University, Ithaca, NY 14853; Department of Ecology and Evolutionary Biology, Cornell University, Ithaca, NY 14853

**Keywords:** Laboratory mice, wild-derived mice, socioecology, space use, social structure, territoriality

## Abstract

Social behavior and social organization have major influences on individual health and fitness. Yet, biomedical research focuses on studying a few genotypes under impoverished social conditions. Understanding how lab conditions have modified social organizations of model organisms, such as lab mice, relative to natural populations is a missing link between socioecology and biomedical science. Using a common garden design, we describe the formation of social structure in the well-studied laboratory mouse strain, C57BL/6J, in replicated mixed-sex populations over 10-day trials compared to control trials with wild-derived outbred house mice in outdoor field enclosures. We focus on three key features of mouse social systems: (i) territory establishment in males, (ii) female social relationships, and (iii) the social networks formed by the populations. Male territorial behaviors were similar but muted in C57 compared to wild-derived mice. Female C57 sharply differed from wild-derived females, showing little social bias toward cage mates and exploring substantially more of the enclosures compared to all other groups. Female behavior consistently generated denser social networks in C57 than in wild-derived mice. The repeatable societies formed under field conditions highlights opportunities to experimentally study the interplay between society and individual biology using model organisms.

## Introduction

Laboratory house mice are the premier model organism in biomedical research due to their small size, rapid breeding cycle, and the ready deployment of precise experimental manipulations using powerful genetic and neurobiological tools^1–6^. The wide availability of classical inbred mouse strains has allowed the scientific community to amass diverse physiological, genomic, neurobiological and behavioral datasets on repeatable genotypes across labs and studies^2, 7–15^. While highly controlled conditions are required for many experiments, there is growing recognition that environmentally impoverished traditional laboratory approaches limit our ability to understand many complex biological processes^16–21^. For example, constrained lab environments inherently limit the study of patterns of space use or social behavior that contribute to social organization in natural populations, yet the consequences of a population’s social organization on individuals is increasingly recognized as a key factor shaping lifetime patterns of health and fitness^22–25^. At the same time, there have been repeated calls to study traditional model organisms, especially mice, under more natural contexts^26–33^. Indeed, studies of lab mice in outdoor enclosures have already revealed effects of more natural conditions relative to traditional laboratory housing on traits including foraging, hippocampal neurogenesis, immunity, microbiome, and cancer progression^34–42^, though we do not yet know the social structure of lab mice under semi-natural field conditions.

Over the last decade, multiple groups have developed high-throughput lab assays using modestly-sized arenas or interconnected cages that allow for increased complexity of social interactions^43–45^. Yet, even relatively large and enriched lab settings^46–48^ fail to capture many of the relevant features of social structures inferred by studies of wild mouse populations to be important to mouse natural history, such as territorial social organization and space use^49–54^. Accumulating evidence suggests that generations of inbreeding and artificial selection in lab mouse strains has impacted their social behavior and interactions in small groups^49^. Indeed, even simple lab behavioral assays can reveal differences in classical lab strain and wild-derived mouse behavior^55–60^. But we have little understanding of the ecological validity of such studies on lab mice because of the lack of studies addressing the socioecology of lab mice under natural or semi-natural field conditions. At stake is not only our understanding of the ecological validity of studies of social behavior in constrained conditions, but also how we interpret and understand the difference between domesticated lab and genetically wild mice more generally.

An immediate solution is to study the social and spatial behavior of lab mice in large naturalistic spaces. Providing ample space for individuals to interact or avoid each other is critical for assessing social structures because it allows animals to freely express their preferences. There is a long history of studies utilizing semi-natural indoor or large outdoor enclosures to study the population biology of house mice under free-range conditions^61–72^. These studies tend to use feral or wild-derived populations of outbred house mice and find that male mice establish and aggressively defend territories occupied by several females and their offspring. Fully adult males are most often associated with high quality territories, while juveniles and subadults typically aggregate in lower quality spaces within the environment^62, 73, 74^. Adult females may compete for nest sites but form strong associations and may even co-nest with close relatives^75, 76^. The competitive environments generated under free-range conditions can have a strong impact on social behavior relative to the lab environment^66, 77, 78^, motivating our study of mice in field enclosures where they can freely compete in a semi-natural environment.

Understanding how lab mice behave under more natural field conditions will inform how their behaviors have evolved relative to wild mice, providing a crucial piece of biological data for the best studied mammalian genotype, C57BL/6J (hereafter ‘C57’). Here we report the space use and social behavior of replicated mixed-sex populations of C57 and wild-derived outbred (hereafter ‘WD’) mice in large outdoor field enclosures. We aimed to address three empirical questions with this study. (1) We sought to determine whether male C57 would establish and defend territories in the field, as opposed to generating societies with a single integrated dominance hierarchy (a common outcome in the lab^48, 79–81)^. We also compare the territorial behaviors of C57 mice and WD outbred mice. (2) We sought to test whether space use and the resulting social relationships that form differ between C57 and WD outbred females. Female fitness in wild mice depends on their abilities to compete for nesting sites^62, 75, 82, 83^ and they show spatial and social biases toward other related females^76^. We hypothesized that lab mouse husbandry practices, which regularly lead to isogenic females living and occasionally breeding at very high density^1^, may have led to changes in the magnitude of social biases towards familiar females. (3) Finally, we sought to describe the social networks that emerged from these individual behaviors, and how differences in behavior between C57 and WD mice scaled up to shape the larger social structure. We hypothesized that, despite the myriad complexities and idiosyncrasies of individual decisions and environmental fluctuations that influence societies, repeated studies of societies that differed in a salient manipulation—in this case the genotype of the individuals—would result in consistently different outcomes across treatments. If this hypothesis is correct, populations with similar initial ecological and demographic conditions will reliably generate similar social structures, suggesting that the biological basis of social organization is amenable to study.

## Results

Over a three-month period (June 2020 -August 2020), we examined the emergent social organization generated in enclosures stocked with adult male (n = 10 per trial) and adult female (n = 10 per trial) C57 (n = 4 trials) and WD outbred house mice (n = 3 trials) over the course 10-day trials. A detailed discussion of the study design, including the density of mice, length of trials, and genotypes tested is provided in the methods section. Each field enclosure (38.1 m x 15.24 m; ∼570 m^2^) contained eight weather protected resource zones arranged in a two by four grid pattern (**Fig. 1**; **Fig. S1A-B**), resulting in a slight excess of mice of each sex relative to the available resource zones. Mice were implanted with PIT tags and activity at the resource zones was monitored via an RFID antenna (see Methods). We obtained high density sampling of mouse RFID reads for all trials (1,307,712 ± 135,646 reads per trial; mean ± SE) and a mean of 6771 ± 275 RFID reads per mouse per. Mice were able to quickly traverse the distance between the zones despite the ground vegetation (minimum inter-zone travel time = 10 seconds; **Fig. S1D**). We estimated individual mouse location for a total of 5653 hours across all trials (mean *=* 808 ± 51 hours/trial). On average, we inferred individual mice spent 4.2 ± 0.1 hours/day at resource zones (range: 1.1 seconds – 18.7 hours).

**Figure 1:**
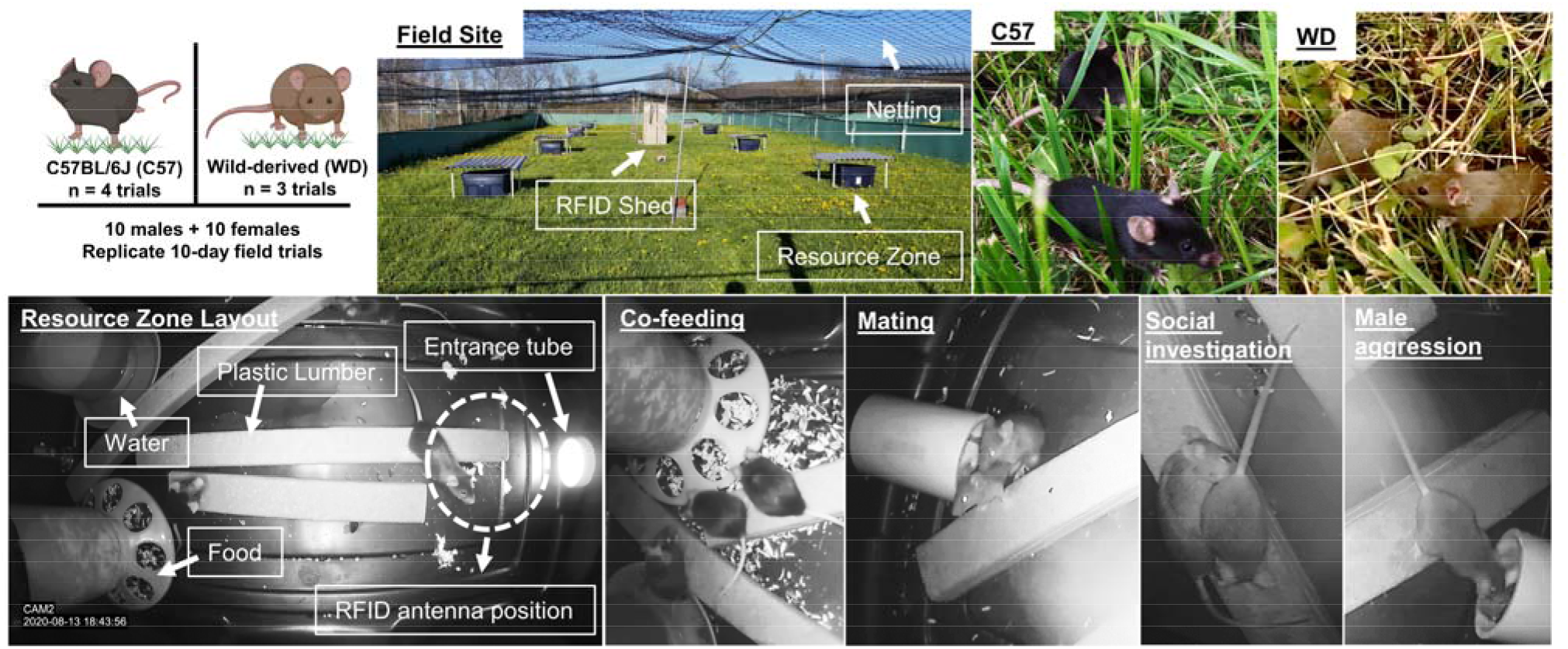
Experimental design for replicate populations of C57 and wild-derived mice in field enclosures. Photos demonstrate the layout of the field enclosures and the eight resource zones arranged in a 2×4 grid pattern. Resource zones had a single entrance tube and food and water towers provisioned *ad libitum.* We observed a variety of behaviors in the resource zones including co-feeding between females, mating and courtship, social investigation, and male-directed aggression towards intruders. Top left image created with BioRender.

### Territory establishment is slower in C57 compared to wild-derived males

Territorial behavior is reported from wild mouse populations and field enclosure studies at moderate density or in lab environments with high physical complexity^64, 73, 84–86^. Territorial males attempt to exclude other males from the spaces that they control, and the ability to compete for and maintain a territory is a key driver of male fitness in freely mating mouse populations^71^. In contrast, in laboratory and natural settings where there is low environmental complexity such that individual dominant mice can readily patrol most of the available space, an alternative social structure where groups of male mice form a social hierarchy predominates^48, 79, 80, 87^. Mouse social hierarchies can vary in the extent of despotism or egalitarianism^48^, though are generally characterized by a dominant alpha who is aggressive to all others and has broad access to space, followed by a linear hierarchy of subordinates^81^. Commensal mouse populations may show a mixture of these social forms, with hierarchies formed among the males residing within a single territory or deme but are nevertheless characterized by restricted movement of mice between territorial spaces.

We sought to determine whether C57 and WD males in our enclosures formed territories or dominance hierarchies. If males form territories in our enclosures, a subset of the males in each trial would each monopolize one or more resource zones to the relative exclusion of others. What’s more, no male would regularly access all resource zones. In contrast, if males establish a dominance hierarchy within the enclosure, the top-ranking male or males are expected to have relatively free reign and regularly access all or nearly all resource zones.

Consistent with previous field enclosure studies of wild-derived mice^73^, we find evidence that males of both genotypes established and defended territories, though the intensity of territorial behaviors was muted in C57 relative to the WD outbred mice. Three related, but distinct analyses support this conclusion.

First, males appear to establish monopolized spaces and rarely visit resource zones that they do not control (**Fig 2A; Fig. S2A**). By the third day, 93% ± 1 of all male-sourced RFID reads within each resource zone belonged to a single male, with zones in WD trials becoming monopolized by males more rapidly than in C57 trials over the 10 day period (*t*_500_ = -2.93, *P* = 0.004; **Fig. 2B**). No individual male consistently accessed all or nearly all resource zones (range of mean daily zones visited = 1.0-3.9, median = 1.6). Instead, males spent the vast majority of their time within their single most visited resource zone, with WD males developing this site fidelity more rapidly than C57 males (*t_67_* = 2.31, *P* = 0.02; **Fig. S2B**). Thus, male space use was consistent with territory defense, and inconsistent with a single integrated dominance hierarchy.

**Figure 2:**
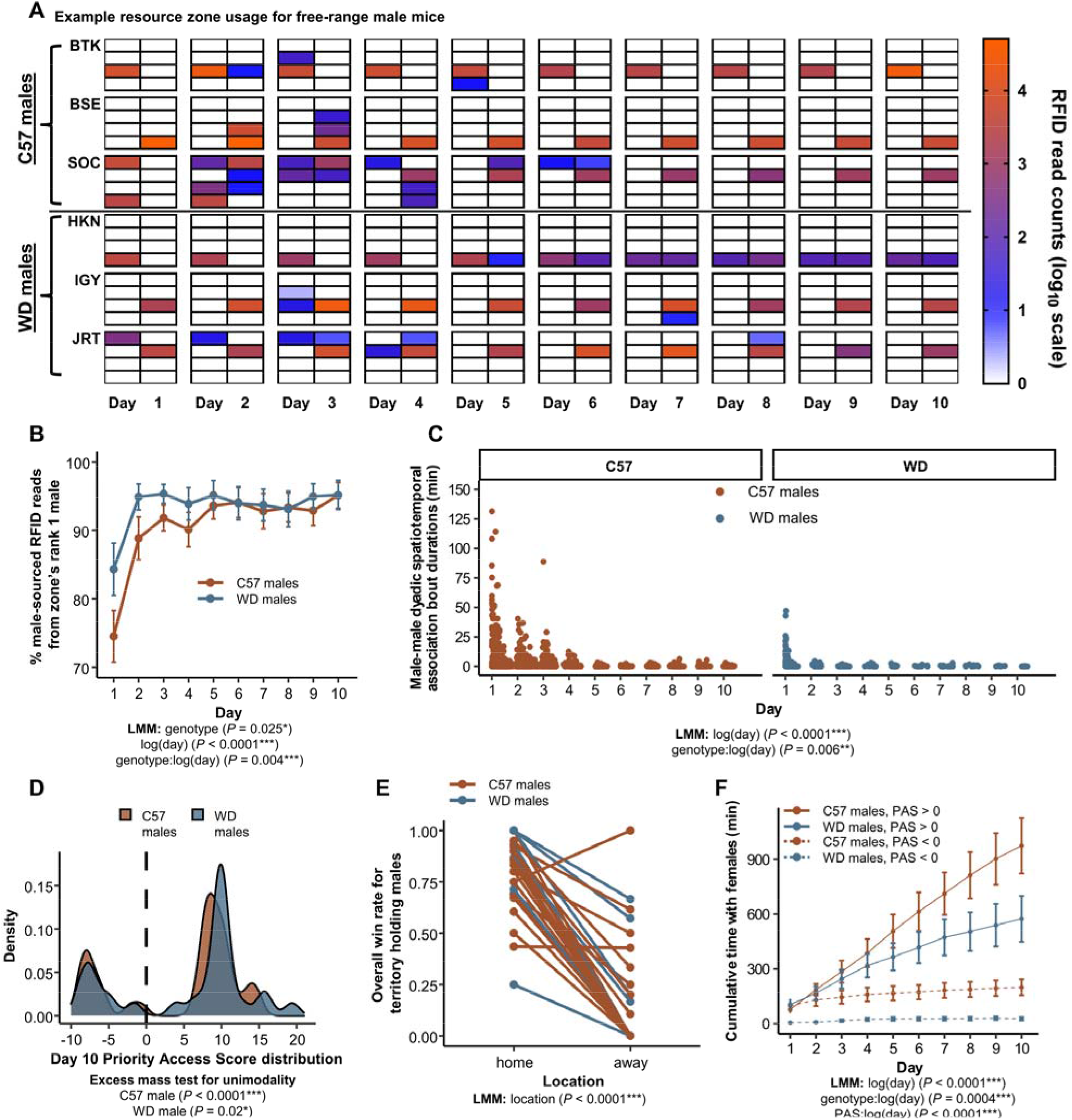
C57 and wild-derived male mice establish territories rather than dominance hierarchies. **(A)** Males generally visited only one or two zones, and no males visited all zones, as shown by representative space use data from three C57 and three WD males. Each 2×4 grid shows the eight resource zones on each of the 10 days of a trial, with redder colors indicating a high concentration of RFID reads. **(B)** WD males form territories more quickly than C57 males. Here the y-axis represents the percent male-sourced RFID reads per zone from the focal zone’s most present male, with values of 100% indicating complete monopolization. **(C)** Male-male dyadic spatiotemporal association bout events were shorter, less frequent, and deteriorated more quickly in WD mice compared to C57 mice. **(D)** To quantify territoriality, we calculated a Priority Access Score, a cumulative metric of resource zone access across the 10 days of the trial. For both WD and C57 males, this territory metric was deviated from unimodality, showing a group of males that had consistent access to resource zones (territory holders) and those without (non-territorial). Here scores near or greater than +10 indicate a male that controlled a resource zone for the duration of the trial. Negative scores are indicative of males that failed to capture a territory. **(E)** Consistent with territoriality, but not a dominance hierarchy, territory-holding males were much more likely to win a spatial dispute with another male, defined as displacing the other male, when the dispute occurred in their home territory as opposed to in a different male’s territory. **(F)** Territorial control (as categorized by having a Day 10 PAS > 0) conferred benefits in the form of increased access to females. This benefit of territoriality was stronger among WD as compared to C57 males (notice that WD males without a territory essentially never spend any time with females, dashed blue line). Data are plotted as means ± s.e.m.

Second, males spend limited time in spatiotemporal overlap with other males. In all trials, males were placed into the enclosure within a resource zone along with their cage mate (brother). Males therefore overlapped in space at the start of the trials, but episodes of male spatiotemporal overlap rapidly decreased in both frequency (*t_58_* = -9.47, *P* < 0.0001, **Fig. S2C**) and duration (*t_2116_*= -13.15, *P* < 0.0001, **Fig. 2C**). The collapse of previously existing cage mate relationships was remarkably rapid in WD males. At least half of all the estimated male dyadic spatiotemporal overlap time in WD mice elapsed in the first 54 minutes of a trial on average (range = 32 – 93 minutes), compared to 590 minutes on average (range = 210 – 1501 minutes; *P* = 0.02) for C57 males. Male-male spatial overlaps shortened in duration over the course of the trial more quickly in WD males as compared to C57 males (*t_2116_* = -2.77, *P* = 0.006, **Fig. 2C**) and were less frequent overall (*t_27_* = -4.14, *P* = 0.0003, **Fig. S2C**), suggesting that the intensity of male-male competition in C57 may be weaker than in genetically wild mice. Strikingly, on days 5-10, when territories were clearly established, the amount of temporal overlap between a dominant territorial male in a given zone and other males that visited that same zone was 86% less than expected if each males’ space use was independent of each other (range across trials = 74% - 97% less; *t_6_* = -58, *P* < 0.0001). This pattern could be explained either by (a) territorial males quickly expelling intruders when they co-occur or (b) non-territorial males biasing their visitation behavior to times that they expect the territorial male to be absent.

To characterize the variation in males’ territorial behavior and resource access, we developed a Priority Access Score (PAS) metric which tracked changes over time in the degree to which mice monopolized access to resource zones relative to same sex conspecifics (see Methods). Briefly, for the 10-day trials reported here, strongly positive final scores (near +10) indicate consistently monopolized a single resource zone on each day of the trial while strongly negative scores (near -10) indicate an individual was consistently excluded from most spaces. High scores (>>10) indicate individuals who monopolized more than 1 zone over the course of the ten-day trial. Scores closer to zero indicate individuals that share spaces to some extent with other individuals of the same sex. Consistent with the hypothesized presence of territorial and non-territorial males we find a bimodal distribution of males with high and low PAS values (excess mass test for unimodality: *P* < 0.02; **Fig. 2D; Fig. S2D;** in contrast, females of both genotypes do not strongly monopolize resource zones at this density, see **Fig. S3B-C**).

The third and ultimate indicator of territory formation rather than a single dominance hierarchy would be for contest outcomes to be predicated on spatial ownership. In other words, for territorial males to win competitive interactions in their territory and lose in other males’ territories such that encounter outcomes depend on location. In contrast, under a social dominance hierarchy, a given male should be win or lose depending on their rank without respect to location. To assess this prediction, we investigated male-male dyadic interactions within territories by identifying when the territory-holding male spatiotemporally overlapped with an intruder. For each such inferred overlap, we identified which animal was the first to leave the interaction (i.e., was displaced and lost acute resource access) and assigned that male as the ‘loser’ of the interaction event. We assigned the male remaining in the zone as the ‘winner’.

We identified 1380 interactions which, as expected, overwhelmingly involved territory holders (n = 1290 events, 93.5% across all trials; mean = 25.6, median = 14, 95% C.I. [15.8, 35.2] per territory holding male). To compare win and loss rates for the same individuals, we examined 32 territorial males that were observed to engage in both ‘home’ and ‘away’ displacement event (n = 23 C57, n = 9 WD; n.b.-many territorial males were not observed in displacement events in an ‘away’ context). We find that overall win rates of territory holders at home (0.82 ± 0.03) are dramatically higher than win rates away (0.18 ± 0.05; *t_31_ =* 12.19, *P* < 0.0001; **Fig. 2E**), with only one male showing a lower win rate at home than away. The one male that had an excess win rate away was only observed to participate in single away displacement event, which he won. More than half of the males (59%, 19/32) lost all their contests when away from their territories. Winning away was rare after the first day of the trial, when boundaries were still in flux; 72% (50/69) of displacement events won away occurred on the first day of the trial. Together, these results support the territory hypothesis and reject a hypothesis where male mice in the enclosures formed a single integrated dominance hierarchy.

Territorial control was associated with increased access to females. Males with positive (>0) final PAS values spent more total time with females than males with negative (<0) final PAS values over the course of the trial (*t_661_* = -3.6, *P* = 0.0004; **Fig. 2F**). The effect of territoriality on males’ access to females was stronger among WD males. Among WD males, those with low PAS values essentially never spent any time with females (range of total time spent with females = 0.73 – 74.5 minutes), even on the first day of the experiment. In contrast, although C57 males with low PAS values spend dramatically less time with females over the course of the experiment as compared to high PAS males, they spend comparable amounts of time with females during the first day of the experiment, suggesting that the competitive exclusion of males from access to females was slower to develop in C57 mice.

Establishment of non-overlapping territories led to similar patterns of space use in both C57 and WD males. Examples of males’ spatial behavior are shown in spatial heatmaps in **Fig. 2A** highlighting the spatial separation and fidelity of males in a single trial (see **Fig. S2A** for full example trials). The males in each trial that were not able to capture a resource zone (those with Priority Access Scores less than 0) appeared to adopt an alternative strategy in which they briefly visited several zones each day. These non-territorial males, who were not tied to a specific space that they needed to defend, tended to visit more zones over time on average as compared to the territorial males (*t_65_* = 2.61, *P* = 0.01; **Fig. S2F**).

### Distinct patterns of space use and social associations in C57 females

Studies of free-living house mice document competition among females for nesting space. Females tend to tolerate their relatives while avoiding or even showing aggression towards unrelated females^76, 82, 83, 88^. At the same time, male territorial structure may shape female behavior as novel males can lead to pregnancy termination through the Bruce effect^89, 90^ and novel males represent an infanticide threat^89–92^. Competition for nesting space and avoidance of non-sire males is likely to be most acute among breeding females^72, 93^. Our data examine how females use space when first introduced into a large novel social environment and assess the extent to which their space-use behaviors are influenced by their social environment.

C57 females exhibited substantial differences in space and movement patterns across several measures compared to WD females as well as both male genotypes (**Fig. 3A; Fig. S3A; Fig. 2A; Fig. S2A**). Unlike WD females and males of both genotypes, C57 female space use was not limited to a few neighboring resource zones, but instead was widespread across the enclosure space. Across sexes and genotypes, mice visited an average of 2.32 ± 0.11 resource zones per day over the course of the trial, though patterns of zone visits varied over time and among individuals. On average, the number of resource zones that animals visited each day increased as the trials progressed (*t_133_* = 13.3, *P* < 0.0001), but this increase was driven entirely by the behavior of C57 females (*P* = 0.52 for non-C57 females; **Fig. 3B**). Although all mice explored an equivalently low number of resource zones during the first several days in the enclosure, by the fourth day C57 females had significantly increased exploration of the available zones compared to all other groups (*P* < 0.05 for daily LMM contrast estimates for Day 4 – 10), which did not differ in their extent of space use.

**Figure 3:**
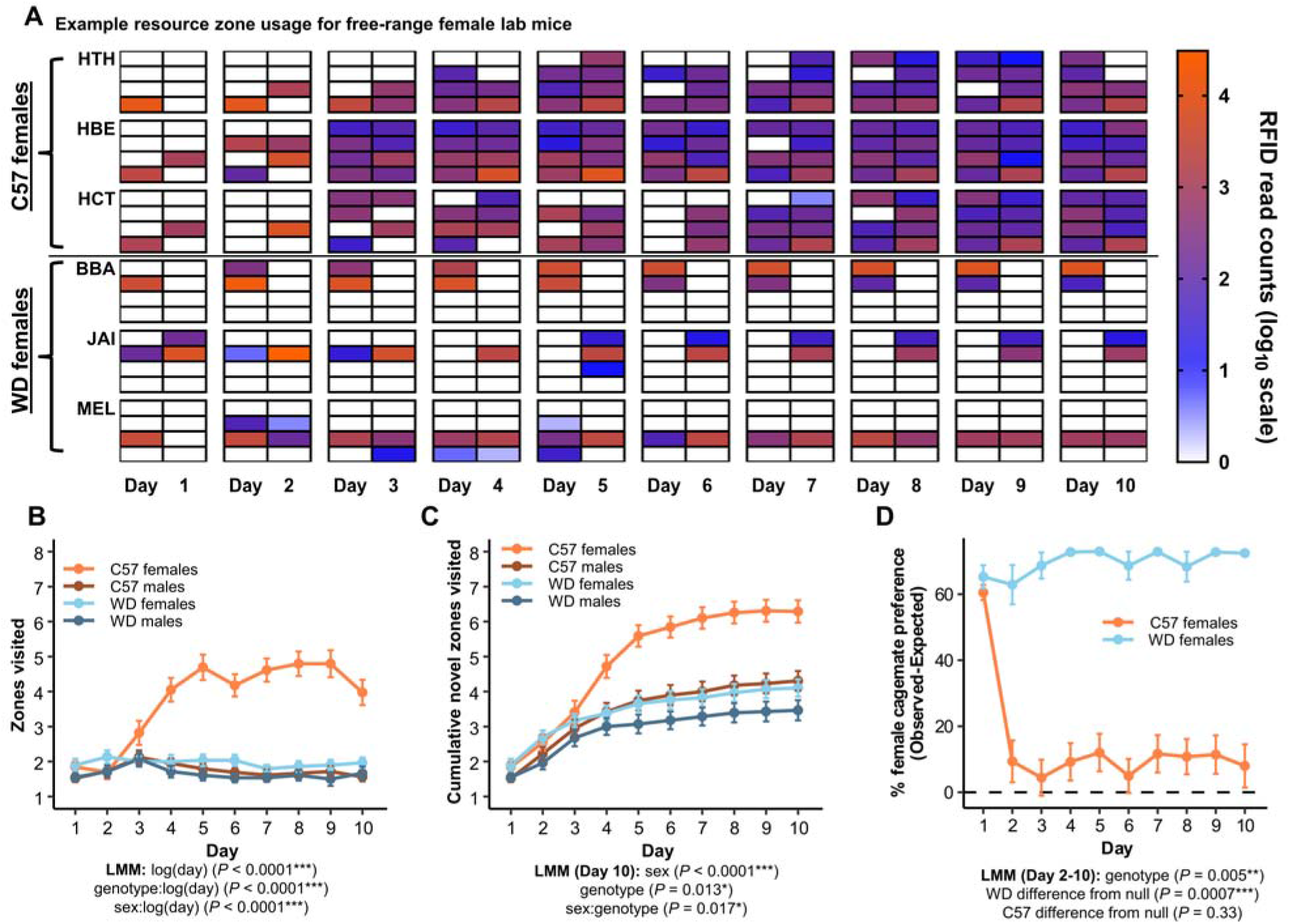
C57 females show distinct patterns of space use and social associations. **(A)** C57 females explore more of the available resource zone spaces than C57 males and WD males and females, as shown by three representative individuals from each strain. Each 2×4 grid shows the eight resource zones on each of the 10 days of a trial, with warmer colors indicating a high frequency of RFID reads. **(B)** All sex and strain combinations visit similar numbers of zones (unique per day) on the first several days of the trial, but C57 females increase the numbers of zones visited relative to any other sex or strain combination as the trial continues. **(C)** C57 females visit more cumulative resource zones over the course of the trial period compared to all other sex and strain combinations. **(D)** WD females show a strong and sustained preference for their cage mates while C57 female preferences for cage mates quickly fall to null expectation levels (as indicated by the dashed line). Note that mice were placed in resource zones with their cage mates on the first day of the trial. Data are plotted as means ± s.e.m.

In addition to visiting more zones on average per day, C57 females visited a greater proportion of all possible zones over the course of the trial (**Fig. 3C**). By the final day of the trial C57 females had visited 6.3 ± 0.4 of the 8 available zones, which is more than C57 males (4.3 ± 0.4; *t_125_* = 5.5, *P <* 0.0001), WD females (4.1 ± 0.5; *t_7.4_* = 3.3, *P* = 0.045), and WD males (3.5 ± 0.5; *t_7.6_* = 4.3, *P* = 0.01). Many more C57 females (42%, 16/38) visited all eight resource zones as compared to C57 males (8%, 3/39), WD females (3%, 1/29), and WD males (4%, 1/28) (generalized LMM: *P* < 0.05 for all comparisons). Finally, C57 females spent less time in their most occupied zone compared to WD females (*t_18_* = 2.3, *P* = 0.04; **Fig. S3E**).

Female house mice exhibit social preferences towards familiar same-sex conspecifics under free-living conditions^76, 82^, but the degree to which this happens in lab strains like C57 is unclear. Our results suggest that C57 females are more tolerant towards females in general and less biased towards familiar social partners. Overall, C57 mice engage in longer female-female spatiotemporal association bouts compared to WD mice over the course of the trial (*t_11910_*= -2.8, *P* = 0.02; **Fig. S3F).** At the same time, the female preference for spending time with cage mates on days 2-10 of the trials was much greater in WD females than in C57 females (*t_5_* = 4.8, *P* = 0.005). WD females (*t_5_*= 7.2, *P* = 0.0007), but not C57 females (*t_5_* = 1.1, *P* = 0.33), differed significantly from the null value based on the prevalence of cage mates within a trial (**Fig. 3D**). Indeed, WD females spent nearly all their female-female social time with cage mates after day 2, whereas C57 females tended to spend less than half of their time with cage mates (**Fig. S3G**). These results suggest that inbreeding and/or selection from lab mouse colony rearing practices may have a profound effect on both female space use and social behavior.

### Repeatable differences in social structure between C57 and wild-derived mouse societies

Analyses of the behavior of each sex show generally similar patterns of space use for males but strongly divergent patterns for females. We next sought to understand how behavioral differences measured for individuals scaled up to larger patterns of social association and the emergent structure of societies.

We first examined how mice overlapped in space and time to determine to what extent individuals associated, as well as the range of spatiotemporal group compositions that arose. For each trial we estimated the time spent in group compositions of differing numbers of males and females (**Fig. 4A**). For this analysis, we considered each mouse’s time separately such that a pair of mice in a zone together for 60 minutes is recorded as two mouse hours.

**Figure 4:**
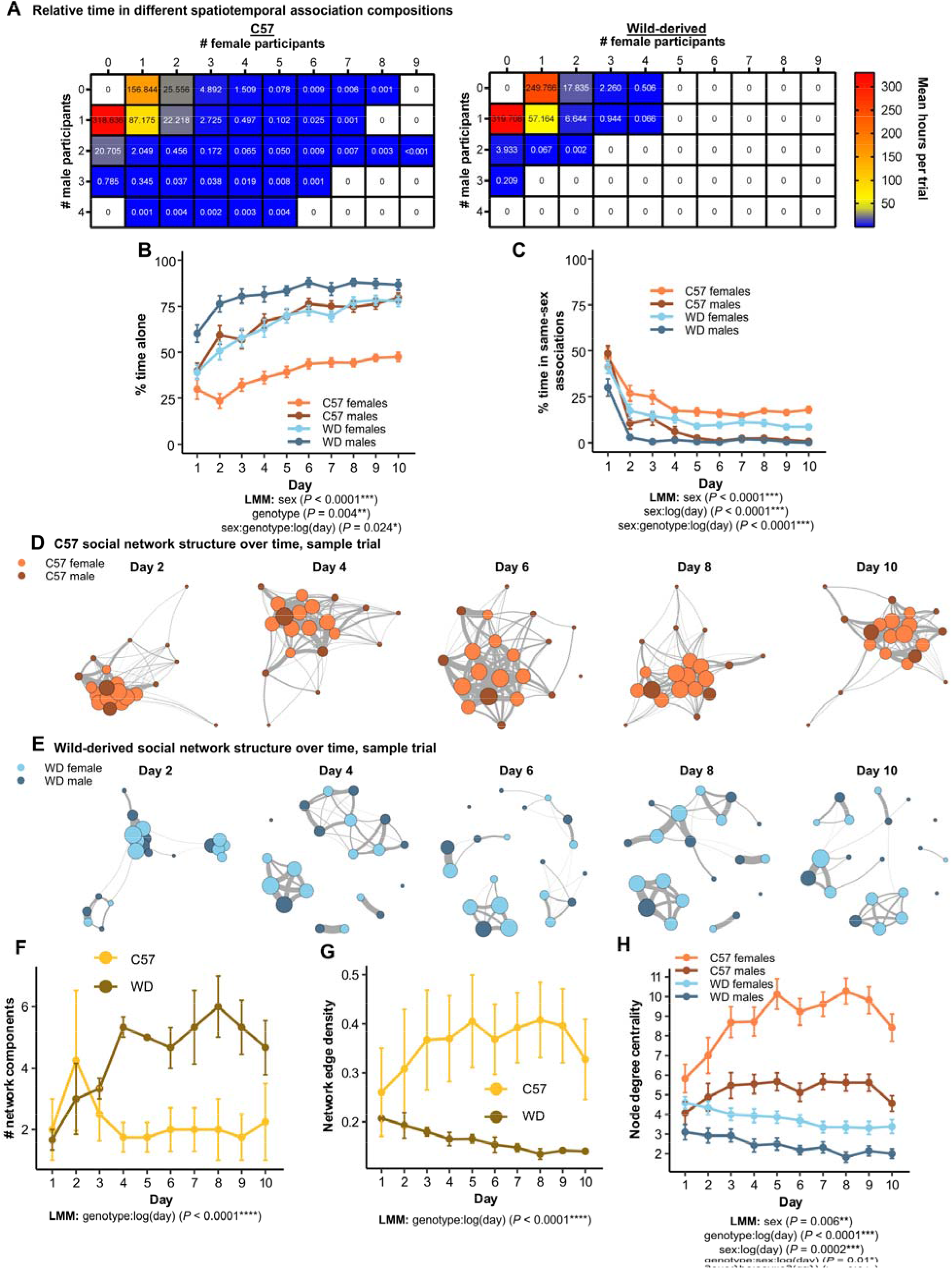
C57 social structures are more interconnected than in wild-derived mouse societies. **(A)** C57 mice spend less time alone and have larger numbers of social participants in association events as compared to WD mice. The contour plot shows the average duration of observed time spent in different male and female group compositions across trials for C57 (left) and WD (right) mice. The numbers reported here are the average number of hours estimated per trial. **(B)** C57 females spend less than half of their total observed time alone throughout the trial, less than any other group. **(C)** All strain and sex combinations decrease the time spent in same-sex groups over the course of the trial, with males of both strains spending minimal amounts of their observation time in same-sex interactions. **(D-E)** Daily social networks from an example C57 trial **(D)** demonstrates a typical pattern of persistently high female interconnectivity while an example outbred trial **(E)** demonstrates increasing network modularity over time. The size of connections between nodes represents the edge weight while the size of nodes reflects the node edge strength, or the sum of all edge weights for a single node. **(F)** Number of network components increases in WD, but not C57, social networks over time. **(G)** Network edge density increases in C57, but not WD, social networks over time. **(H)** C57 females have consistently higher measures of node degree centrality, indicating that they are consistently meeting large numbers of social partners on each day of the trial, relative to other sex and strain combinations. Data are plotted as means ± s.e.m.

In each trial, mice were alone for most of the time that they were recorded at resource zones (range 50-64% for C57; 70-82% for WD mice, solitary mouse time per trial; *P* = 0.03 based on the probability of all four C57 trials having the lowest values; **Fig. 4A)**, but the proportion of time that individuals spent alone was strongly predicted by sex and genotype. On average, males spent a greater proportion of recorded time at resource zones alone than females (*t_1339_* = 5.4, *P* < 0.0001; **Fig. 4B**). Overall, WD males increased their time alone over the course of the trial (*t_1339_* = -3.1, *P* = 0.002; **Fig. 4B**) and were especially likely to spend time alone compared to all other groups; all of them (29/29) spent more than 50% of their total recorded time alone. In comparison 83% of WD females (25/30), 78% of C57 males (31/40), and only 18% of C57 females (7/40) spent most of their recorded time alone. Given the interest in the biology of social isolation in mice^94–98^, it is notable that when given the opportunity to freely interact, many mice spent a significant portion of their observed time alone over the course of their trial.

Though individuals spend a large portion of their time at the resource zones by themselves, we estimated more than 2000 mouse hours of spatiotemporal associations across the seven trials, defined as time with two or more mice at the same zone simultaneously (**Fig. S1F**). Dyadic interactions accounted for >50% of estimated association time in both genotypes (62-80% in C57, 78-86% in WD), though larger aggregations of mice were detected in all trials (**Fig. 4A**). On average, females spent a greater portion of their recorded time in same-sex associations than males over the course of the trial (*t_1339_* = -6.5, *P* < 0.0001; **Fig. S4A**). Most mice (77%, n = 103/134) spent >50% of their recorded association time in opposite-sex groups, with males increasing this metric over the course of the trial relative to females (*t_1165_* = 6.6, *P* < 0.0001; **Fig. S4B**).

To investigate the emergent group structure of both genotypes, we analyzed the total and daily social networks formed for each trial. Overall, C57 mice formed more connected networks than WD mice, a difference which was largely driven by high levels of C57 female sociability (**Fig. 4D-E**). WD networks increased in the number of graph components – the portions of the network disconnected from each other – over time (*t_61_* = 4.86, *P* < 0.0001; **Fig. 4F**), echoing the demic structure reported for many wild mouse populations^53, 99, 100^. In contrast, C57 networks became more connected over the course of the trials. Over time, the network edge density – a measure of the proportion of edges actually observed out of all possible edges in the network – increased in C57 social networks, but not in WD networks (*t_61_* = -5.4, *P* < 0.0001; **Fig. 4G**). Notably, the two genotypes show completely non-overlapping distributions for network edge density from days 4-10 (*P* = 0.03, non-parametric, each day). We note here that the differences between genotypes reported in this section are based on whole network level metrics indicating that the basic overall structure of social organization differs between the genotypes.

Females of both genotypes had high degree centrality measures compared to their respective males, indicating females encountered more unique social partners on each day of the trial. There was a significant three-way interaction between sex, genotype, and time, such that C57 females rapidly increased their network centrality measures compared to all other sex and genotype combinations (*t_134_* = 2.6, *P* = 0.01; **Fig. 4H**). Thus, many of the differences we see in social networks between the genotypes is driven by the propensity of C57 females to co-occur with many distinct individuals. By the final day of the trial, C57 mice had overlapped with many more of the available social partners present in the enclosures as compared to WD mice (*t_5.48_* = -3.9, *P* = 0.0098; **Fig. S4C**), who failed, on average, to ever meet more than 50% of the potential social partners, at least within or around resource zones. Intriguingly, females of both genotypes showed high levels of vertex page rank scores, indicating that females of both strains maintain higher levels of connectedness over time than their respective males (*t*_1343_ = 3.2, *P* = 0.001; **Fig. S4D**).

## Discussion

Our replicated semi-natural field enclosure experiments demonstrate that C57BL/6J lab mice broadly recapitulate many of the behaviors of wild-derived outbred mice in free-ranging conditions, including clear evidence of male territories, but have different emergent social structures. This difference is largely due to C57 females being more exploratory and showing less biased patterns of social association. The organization of mammal societies is influenced by ecological^101–103^, demographic^104–, 107^, and phylogenetic factors^108–110^, each of which were controlled for in our trials. Thus, these data show that genotype can have a strong effect on social structures in mammals^111, 112^. In this case, it demonstrates key social behaviors that have been altered during the process of lab mouse domestication^49^. These data also highlight the flexibility of mouse social behaviors across diverse ecological and demographic conditions. For example, in contrast to lab studies at high density, which identify dominance hierarchies among both lab and wild-derived male mice^47–49, 113^, the males in our lower density populations consistently formed and defended territories regardless of genotype (**Fig. 2**).

While our experiment only examined one set of ecological and demographic conditions, it demonstrates a tractable field approach in which variables including food resources, defensibility of spaces, and demographic compositions are all easily tunable. There has been considerable interest in recent years in high-throughput measures of mouse social behavior in lab settings^43, 45–47, 114^ and our study provides a blueprint for similar studies to be conducted under semi-natural outdoor settings using different socio-ecological conditions or genotypes. Studying reproducible genotypes of mice under similar settings in other locations and seasons provides an exciting avenue to examine the role of environmental variation in shaping individual phenotypes and population dynamics.

What drives the difference that we saw in female space use across our trials (**Fig. 3**)? Space use in female mammals is often predicted by intra-sexual competition for food resources and nest sites^115^, but resource availability and population density were identical across trials in our study. This suggests two non-mutually exclusive possibilities. First, there may be a genetic difference shaping space-use behavior between C57 females and wild-derived outbred females. For example, this could be manifest as an increased tendency to explore irrespective of their social environment. Alternatively, female behavior may respond to the social conditions present in our trials, which differ in some key respects between the two genotypes. C57 females experience a society where everyone has very high genetic similarity, while WD females experience a world with variation in relatedness. It is likely that mouse behavior in general might be sensitive to these parameters, though studies of social interactions in the lab frequently utilize single inbred strains of mice^116–118^. Female mice respond to variation in perceived relatedness between themselves and males^69, 119, 120^, and thus it may be the case that females explore more in conditions when all social partners are genetically homogeneous but show more restricted space use when social partners are genetically heterogenous. In wild house mice, infanticide risk from both male and female conspecifics is thought to be a major driver of social behavior in females^121–123^. As a result, wild female house mice will aggressively defend space from other females^72, 82, 83, 124, 125^. In contrast, C57 mice have been bred to live in cages at high densities, especially among females, and this is associated with lower female aggression compared to wild-derived mouse genotypes^126^. Differences in relative tolerance of other females may be a key driver of the observed differences in social organization between C57 and outbred females in this study. Understanding how innate behavioral differences among genotypes and emergent properties generated by social interactions shape mammalian societies is an exciting future direction.

Male space use in rodents and other mammals is frequently linked to patterns of female space use^115, 123^. Yet, despite differing patterns of female space use between genotypes, the male spatial and social structures across genotypes were quite similar, highlighting that some aspects of social organization are relatively less sensitive to other features of a population’s socioecology. Perhaps one of the most striking features of our study is the speed with which male-male social interactions decrease in frequency, especially among wild-derived outbred males (**Fig. 2C**). Previous studies of wild mouse behavior have reported males will defend territories and attempt to exclude other males^61, 65^ and our data show this behavior is retained in male C57 mice. The formation and physiological consequences of dominance hierarchies among male mice have been the subject of recent study in the lab^48, 49, 113, 127^, but our results suggest that when given ample and defensible spaces male mice will tend to avoid interacting with others and form individual territories rather than a single integrated dominance hierarchy. Differences between territoriality and dominance behaviors remain poorly understood at the mechanistic level. Social dominance hierarchies and the establishment of territorial boundaries could be mediated by the same physiological and neurological mechanisms, or they might be mediated by distinct mechanisms. The process by which dominant males recognize known, tolerated subordinates may be fundamentally distinct from the process by which a territorial male characterizes intruders as well as neighbors versus strangers^128, 129^.

Across repeated trials, we identified differences in the higher-level social organizations of C57 lab mice and their wild-derived outbred counterparts (**Fig. 4**). Studies of social structures tend to come from idiosyncratic populations living in the wild, meaning that studies of social behavior in natural conditions are rarely replicated^130–132^. Studies of free-living populations are critically important, but this non-replicability makes understanding the specific genetic, neurobiological, ecological, and demographic factors influencing complex behavior challenging. The repeatability of social organization demonstrated here suggests that future work manipulating aspects of physiology or neural function in free-range mice will offer a unique opportunity to study not just differences in individual behavior, but also how and whether specific behaviors reliably influence society.

## Methods

### Animals

We examined two genotypes of *M. m. domesticus,* C57BL/6J (C57) and wild-derived (WD) outbred mice. We initially obtained C57 (#000664) mice from The Jackson Laboratory (Bar Harbor, Maine, USA). We maintained a wild-derived outbred mouse stock from intercrossing 6 different wild-caught families generated through distinct initial pairings of wild mice from Saratoga Springs, NY, USA, trapped by MJS in 2013^133^. These mice are descended from the same initial collection in Saratoga Springs that gave rise to the wild-derived inbred mouse strains SarA/NachJ (#035346), SarB/NachJ (#035347), and SarC/NachJ (#035348) available from the Jackson Laboratory. All of the mice used in this study were bred in our lab colony, under standard housing conditions. All mice were between 15 and 28 weeks of age when they were released into the field enclosures.

Rather than merely qualitatively comparing our data to published data on mouse behavior in field enclosures or semi-natural indoor settings^52, 64, 73, 137, 138^, we conducted identical, simultaneous trials using wild-derived outbred mice to allow for a direct quantitative comparison of data collected using the same methods and the same physical environmental conditions. We note that while lab mouse ancestry includes three house mouse subspecies, the C57 genome is >90% *Mus musculus domesticus*^139^. Therefore, as a ‘control’ wild mouse population, we used outbred mice generated from multiple *Mus musculus domesticus* lines derived from wild house mice caught in upstate New York. Thus, the comparison group of mice were born and raised in the lab for multiple generations but have a wild-derived outbred genetic background. The population density in our enclosure was 0.034 mice/m^2^, which falls within the range of typical population densities reported for wild mice^140^ (∼0.0011 – 0.11 mice/m^2^) and is the same order of magnitude as initial populations in other field enclosure studies of wild-derived mouse behavior^64, 69^.

### Ethical note

All experimental procedures adhered to guidelines established by the U.S. National Institutes of Health and the ASAB/ABS guidelines for the use of animals in research^134^ and have been approved by the Cornell University Institutional Animal Care and Use committee (IACUC: Protocol #2015-0060). All animals were briefly anesthetized and implanted with dual PIT tags (12mm) prior to introduction to the field enclosures. This procedure is minimally invasive and is consistent with recommendations by the veterinary and animal care staff at Cornell University. PIT tag weight is negligible, and anesthesia removed any discomfort associated with handling and implantation of the tags. We observed no changes in mouse behavior as a result of this procedure, as assessed by normal food and water intake and daily activity on home cage running wheels.

After the 10-day observation period, >50 live-catch traps (H.B. Sherman, Tallahassee, FL, USA) were baited with nesting material, sunflower seeds and a moistened cotton ball and placed in a grid pattern in the enclosures in the evening (1900-2200 hours) and were checked for occupancy the following morning (0600-0900 hours). Trapping continued until all the mice were recovered or identified as deceased or missing (a conclusion reached if there were no RFID reads in the enclosure for a 24-hour period after 3 days of trapping; see Supplementary Material). The trap locations were recorded, and the individual identities of the mice were confirmed using a handheld RFID reader (BioMark, HPR Lite). Mice were immediately transported to a clean cage with ample nesting material, sunflower seeds and water. At the conclusion of the experiment all mice were euthanized using carbon dioxide inhalation followed by decapitation for the collection of tissues for future analyses.

### Study design

Field studies of wild populations provide a powerful means to link aspects of organismal biology to selection, but are typically hampered by a lack of replication^130, 135^. Enclosure studies done over short, but biologically relevant, time periods provide an opportunity to observe replicate populations across multiple trials. A fundamental feature of enclosure studies, as compared to wild populations, is that the experimenters must choose among a range of possible resource distributions, population densities, sex ratios, etc. as the starting conditions of the trials. House mice naturally live under a wide range of resource distributions and densities across non-commensal and commensal settings^62, 67, 74^, meaning that there exists a large set of naturalistic conditions rather than a single most appropriate condition for studying mice. While studying social organization across a range of socio-ecological starting conditions will be informative, for this study we chose a single common set of initial resource distribution, density, and sex ratio conditions for all trials. For our trials, we released 10 males and 10 females into semi-natural field enclosures with eight resource zones which we monitored using a commercially available RFID system. This experimental design allowed us to assess the consistency of social behaviors and structures formed within the enclosures across replicates and to isolate the effect of host genetic background on these outcomes.

To initiate each trial, we placed mice implanted with passive integrated transponder (PIT) tags (see RFID analyses section) into one of eight resource zones (see Field site section) with their same-sex sibling cage mates in the evening shortly before sunset, meaning that all individuals started the trials at a resource zone with one or more familiar social partners (see Supplementary Material for details on cage mate and relatedness information). We allowed trials to proceed for 10 days. This trial length was chosen because it allowed us to avoid females giving birth in the enclosures and made it feasible to conduct multiple replicated trials in the same enclosures over the course of a single summer. House mouse social structure has been reported to vary seasonally and with shifts in demographic parameters in natural populations^50, 136, 137^, so we note that different conditions or studies conducted at other times of year may yield different results. The experiment reported here focuses on behavior during the initial formation and establishment of social structures within replicated mouse populations in our enclosures while controlling for seasonal and demographic variation. Over the course of 10 days, mice explored the enclosures and resource zones and engaged in a variety of social interactions with conspecifics including courtship, mating, co-nesting, and fighting (**Fig. 1**; **Video S1**). As our goal was to test hypotheses regarding patterns of mouse space use and social structure, we focused our analyses on the large RFID dataset.

### Field site

We conducted field work at Cornell University’s Liddell Laboratory Field Station in Dryden, New York, USA using two adjacent and identically sized outdoor field enclosures. Our enclosures are approximately 9,000 times larger than the area of a standard laboratory mouse cage (**Fig. S1B**). The walls of the enclosures were made from sheet metal and stood approximately 1.2 m tall and extended 1.2 m into the ground to prevent the mice from tunneling and moving between the enclosures. Each enclosure was covered with netting to prevent aerial predation, and gravel was spread along the interior perimeter of each enclosure to discourage digging near the walls. Three days prior to releasing mice into the enclosures, we trapped in and around the enclosures to capture and remove any small mammals or snakes from the enclosure. The enclosures contained a mixture of local perennial grasses and plant communities which were mowed to a height of ∼5 cm prior at the start of each trial to maintain similar ecological starting conditions across trials.

We supplied all resource zones with food and water accessible by the mice *ad libitum.* Resource zones were covered with waterproof corrugated roofing material attached to a polyvinyl chloride (PVC, 1” conduit, Home Depot, USA) frame to shade the zones during the day and provide protection from rain. Resource zones were comprised of two nested 32-gallon storage tubs (Rubbermaid, USA) and a single PVC entrance tube (50 mm diameter) through which the mice could freely enter or exit the zone. Each resource zone contained feeder towers containing food and water in excess (∼50 grams of mixed sunflower and bird seed, and 2 liters of water). Several pieces of plastic lumber were added to provide edges and elevated locations for the mice to perch within the resource zones.

At least 24 hours prior to release in the enclosures, all subjects were placed into a stereotaxic frame (Kopf Instruments, Tuhunga, CA, USA) and briefly anesthetized with isoflurane (3-5%). Mice were subcutaneously implanted with dual PIT tags (BioMark, Boise, ID, USA) in the dorsal flank and periscapular region. Each resource zone was equipped with a 15 cm RFID antenna connected to a centralized data acquisition unit (Biomark, Small Scale System, Boise, ID, USA). Antennas were placed directly beneath the floor adjacent to the PVC zone entrance tubes to increase the likelihood of capturing mouse activity during entrances and exits from the resource zone. Scanning for PIT tags within the antenna range occurred at approximately 2-3 Hz continuously for 10 days.

### RFID analyses

We monitored the resource zones continuously over the trial period via a RFID antenna placed beneath the sole entrance into the zone (**Fig. 1**; **Fig. S1C**). Since the RFID antenna detected tags both above and below the horizontal plane of the antenna and mice moved underneath the resource zones, we inferred RFID reads to indicate that mice were within a space inclusive of the inside and underside of the storage tubs.

To convert instantaneous RFID reads into estimates of how long mice spent at the resource zones, we grouped RFID reads into discreet resource zone visits with estimated durations (**Fig. S1E-F**). As expected, the total number of visits to a zone strongly correlated with the total estimated duration of time spent at a zone (Spearman’s correlation, *R* > 0.84 for all genotype and sex combinations; **Fig. S1G**).

We conducted two types of RFID analyses focusing on (1) when and at what zone each mouse was detected and (2) estimating how long they spent at each zone. In the first set of analyses, we examined whether or not mice visited a given zone during each day of the trial (**Fig. 2A**; **Fig. 3A-C; Fig. S1G; Fig. S2A&F; Fig. S3A**). We estimated the minimum distance travelled using consecutive transitions in the RFID reads between antennas in different resource zones and the known spatial layout of the enclosure (**Fig. S3D**). To assess the degree to which resource zones were exclusively accessed by individual mice, we examined the percent of same-sex sourced RFID reads in zones for each mouse (**Fig. 2B**).

In our second set of analyses estimating the durations of time mice spent in zones, we first examined the time elapsed between consecutive RFID read events for each mouse within each resource zone (the RFID inter-read interval). We found that the distribution of all RFID inter-read intervals was heavily skewed (min = 1s, median = 1s, mean = 16.4s, max = 32,683s; **Fig. S1E**). We grouped RFID reads for each mouse for each zone into visitation bouts using a 139 second (the cut-off for capturing 99% of all the within-mouse inter-read interval values within a single zone) sliding window method such that an RFID read which occurred within 139 seconds of the previous read extended the visit bout duration up to that read. Transitions between zones automatically ended and started bouts in the first and second zones, respectively. We assigned isolated single RFID reads (with no other reads within 139 seconds before or after the focal read) a visit bout duration of 1 second. Using this dataset, we estimated the time mice spent in resource zones (**Fig. S2B**). Mice were defined as participating in a spatiotemporal association bout when two or more mice had overlapping resource zone visit bouts within the same zone (see **Fig. S1F** for a graphical schematic of the approach). We used the spatiotemporal association bout dataset to estimate the duration and frequency of male-male (**Fig. 2C**), female-male (**Fig. S2E**), and female-female (**Fig. S3F**) dyadic association bouts. We omitted a subset of animals from a subset of days for all spatial and social analyses when they received no reads over multiple days and were presumed dead (see Supplementary Material).

### Priority Access Score calculation

Priority access scores were calculated separately for male and female mice within a trial. First, we calculated the time a given mouse (*M*) occupied a resource zone (*Z*) as a percentage of the total time that zone was occupied by same-sex conspecifics on a given day (*D*).

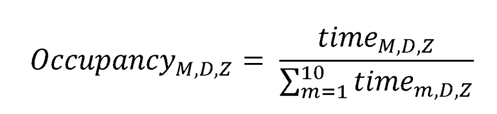

Next, we calculated a daily Capture Score by summing the Occupancy values for all available zones. Mice that did not have an occupancy value of greater than 0.5 (in other words, a majority share of the time spent in any zone), were penalized by subtracting 1 from the final Capture Score. The penalty indicates that on a given day, a mouse failed to capture any of the zones that mouse visited.

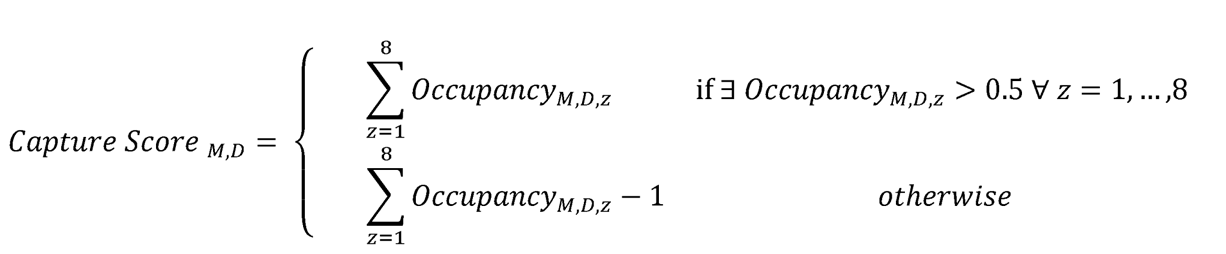

To see how access to zones changed over time, we took the cumulative sum of an individual’s Capture Score ordinally across each day of the trial to derive an evolving Priority Access Score on each day of the trial.

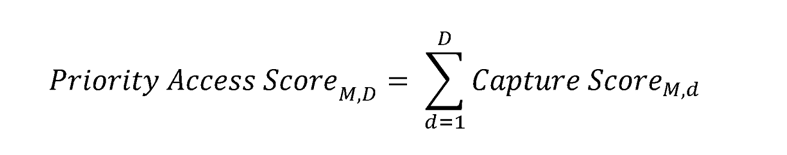

As an example, if one male (male A) occupied a single resource zone every day of the trial for 4 hours a day, while another male (male B) accessed only that same zone for 1 hour per day, and the zone was visited by no other mice, male A’s daily Capture Score would equal 0.8, (because he controlled 4 out of 5 hours), while male B’s daily Capture score would equal -0.8 (because he controlled 1 out of 5 hours and received a one-point penalty for not controlling any zones). If this pattern of visitation remained unchanged for all 10 days, then male A’s final Priority Access Score would equal 8, while male B’s Priority Access Score would equal -8. The Priority Access Score value thus provides a temporally evolving measure that captures the dynamics of territory formation, maintenance, and collapse (**Fig. S2D; Fig. S3B**).

### Male spatiotemporal overlap and win/loss at home and away

We used the zone visit bout and spatiotemporal association bout dataset to investigate the degree to which territorial and intruder males avoided spatiotemporal overlap. Across all zones for the last six days of the trial, we independently calculated the total time territorial and intruder males spent in zones and the amount of that time that was spent in spatiotemporal overlap. We derived null expected values of territorial and intruder male spatiotemporal overlap ((total territorial male hours in zones * total intruder male hours in zones) / (24 hours * 6 days * 8 zones)) and examined the percent difference between the null expectation and the observed time territorial and intruder males overlapped. We chose the last six days of the trials for this analysis as territorial relationships were clearly established by this point in the trial.

Next, we used the RFID and spatiotemporal association bout data to estimate the dynamics of male spatiotemporal overlap and territorial intrusion. We inferred contests by identifying instances in the dataset where a single male within a zone was joined by an additional male, followed by one of the males leaving. We identified the male participants as either the territory holder of that zone or as an intruder based on whether the focal male had captured >50% of all of the male-derived RFID reads within the zone during the last 24 hours. We derived ‘win’ and ‘loss’ rates both within and away from a home territory for each male participant on each day of the trial by assigning the remaining male and the male that left as the ‘winner’ and ‘loser’ of the contest, respectively (**Fig. 2**).

### Female spatiotemporal overlap and cagemate associations

Using the known cage mate relationships between individual females, we examined the total time female mice spent in spatiotemporal association with cage mate (sibling) versus non-cage mate females (**Fig. S3G**). Based on the number of cage mates and non-cage mate female social partners available on each day of the trial (which fluctuated for some trials based on mice disappearing from the trials, see Supplementary Material), we calculated a daily null value of the expected time spent with cage mates and the percent observed value deviation from the expected separately for each female (**Fig. 3D**).

### Social network analyses

We used the spatiotemporal association bout data set arranged in a group by individual matrix to construct social networks. Daily weighted adjacency matrices were derived from a Simple Ratio Index calculation based on counts of binary participation in spatiotemporally overlapping mouse zone visit bout events (where zero indicates that individuals were never in the same spatiotemporal association bout and one indicates that individuals shared all spatiotemporal association bouts**; Fig. S1F**) using the *asnipe*^141^ package in R 4.1.2 (R Development Core Team). All networks were constructed using the weighted matrices and the *igraph*^142^ package in R. Node sizes in our network reflect the node edge strength, or the sum of all connected edges to the node, while edge widths reflect the edge weight derived from the daily weighted adjacency matrices. If an animal died or went missing, we no longer included it in a given trial’s daily network^143, 144^ (see Supplementary Material).

### Statistical analyses

We built mixed effects models using R 4.1.2 (R Development Core Team) and the R packages *lme4*^145^, *lmerTest*^146^, and *emmeans*^147^ to examine relationships between predictor and response variables. We include the full statistical tests and model outputs for all analyses in the Supplementary Material. Most analyses are conducted with a repeated measures design where data is examined for each day per mouse or other relevant unit as appropriate using mixed effects models. Across models we considered random effects of trial, mouse, and/or time in the trial. We included relevant random intercepts and random slopes in our models as appropriate – random slopes of time were generally included, but we occasionally needed to simplify random effects structures when necessary to avoid singular fits. All model formulae, including random effects structures, are explicitly reported in the Supplemental Material. For testing for unimodality, we used the *multimode*^148^ package with “ACR” method^149^. Graphing was done in R using the package *ggplot2*^150^ and in GraphPad Prism 9.3 (www.graphpad.com). We report all means ± standard error (SE), unless otherwise stated, and consider all values statistically significant when *P* < 0.05.

## Declarations

### Consent for publication

All authors have read and approved this manuscript for publication.

## Acknowledegments

We thank Melissa R. Warden for helpful discussions during the planning of the experiment. We thank Gary Olz and Russel L. Ligon for assistance in preparing the field site. Funding was provided by Cornell University (Neurobiology and Behavior Departmental Grant to CCV), the USDA (Hatch Grant, NYC-191428 to MJS), NIH Animal Models for the Social Dimensions of Health and Aging Research Network Pilot Grant (NIH/NIH R24 AG065172 to MJS), and the NIH (R35 GM138284 to AHM).

## Author contributions

Conceptualization: CCV, MJS. Study Design: CCV, MJS. Methodology Design: CCV, MJS. Field Work: CCV, DDS, CHM, MSC, AJM, MJS. Data Curation & Code: CCV. Data Analysis: CCV, MNZ, SXH, MKA, AMG, LCM, MJS. Figure Creation: CCV, MJS. Writing – Original Draft: CCV, MJS. Writing – Review & Editing: CCV, MNZ, AJM, MJS. Supervision: CCV, AJM, MJS. Funding Acquisition: CCV, AJM, MJS.

## Code and data availability

All statistical output tables are included as supplementary files. Code and scripts for reproducing figures and analyses are available on GitHub (https://github.com/calebvogt/2020_LID). All data files are available on Zenodo (https://doi.org/10.5281/zenodo.6425497).

---

**Figure S1:**
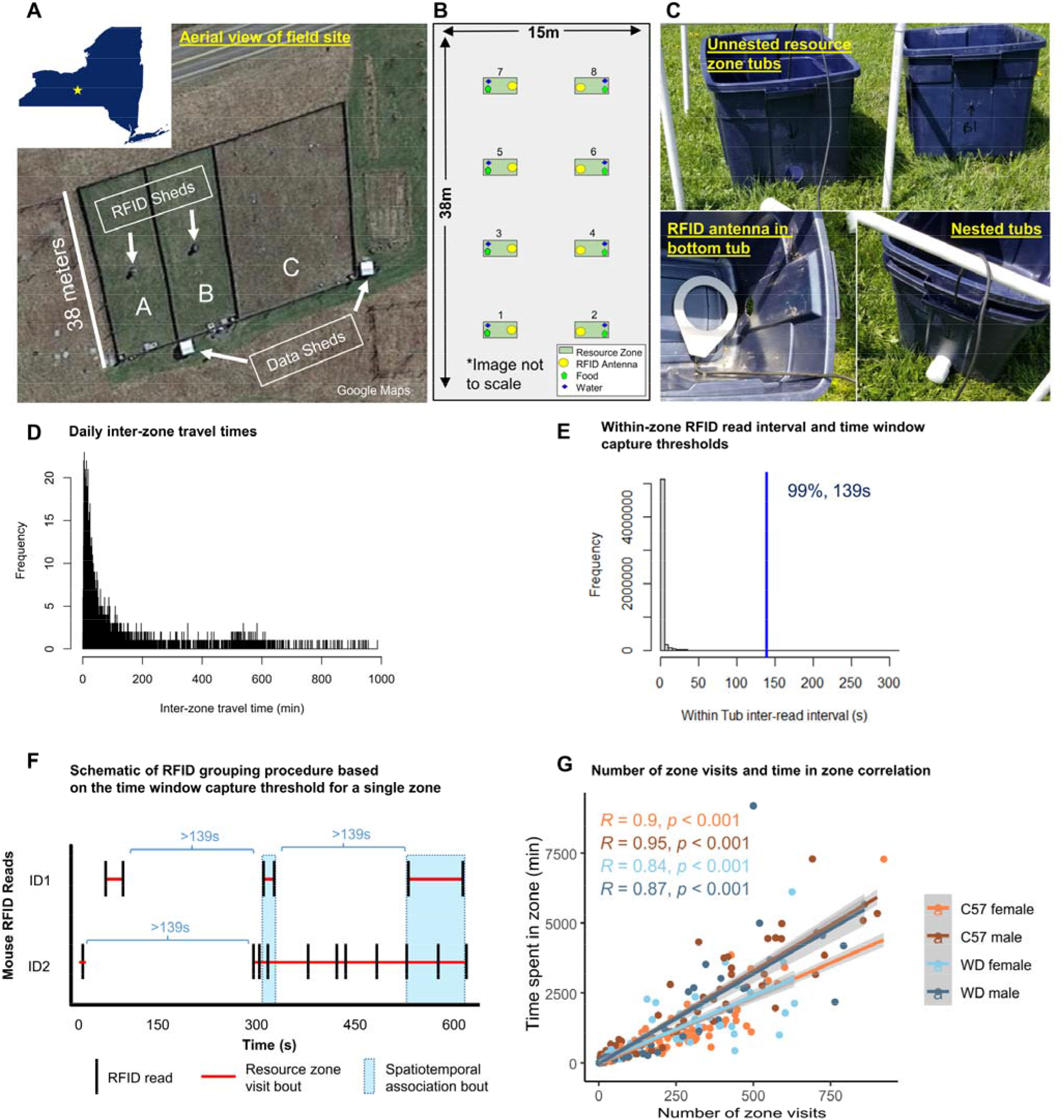
Field site setup and RFID duration bout window selection. **(A)** Satellite image of the field enclosures showing the position of the data sheds housing the computer for downloading RFID data from the central RFID sheds. **(B)** Graphical schematic of the Alpha and Bravo enclosures indicating the resource zone layouts. **(C)** RFID monitoring of the resource zones. Two storage tubs were nested with a RFID antenna placed between them, beneath the entrance tunnel to prevent mice from directly contacting the antenna and wire. **(D)** Histogram of the daily inter-zone travel times for all mice for all days. **(E)** Histogram of the within zone inter-RFID read intervals and the 139 second threshold capturing 99% of all inter-RFID read intervals within the same zone which was used to group RFID reads into resource zone visitation bouts (see Methods). **(F)** Schematic of the time window capture threshold grouping procedure to determine resource zone visit bouts and spatiotemporal association bouts using mock RFID data and the time window capture threshold (139 s). **(G)** Correlation of estimated duration spent in each zone and the number of visits to that zone for all sex and strain categories.

**Figure S2:**
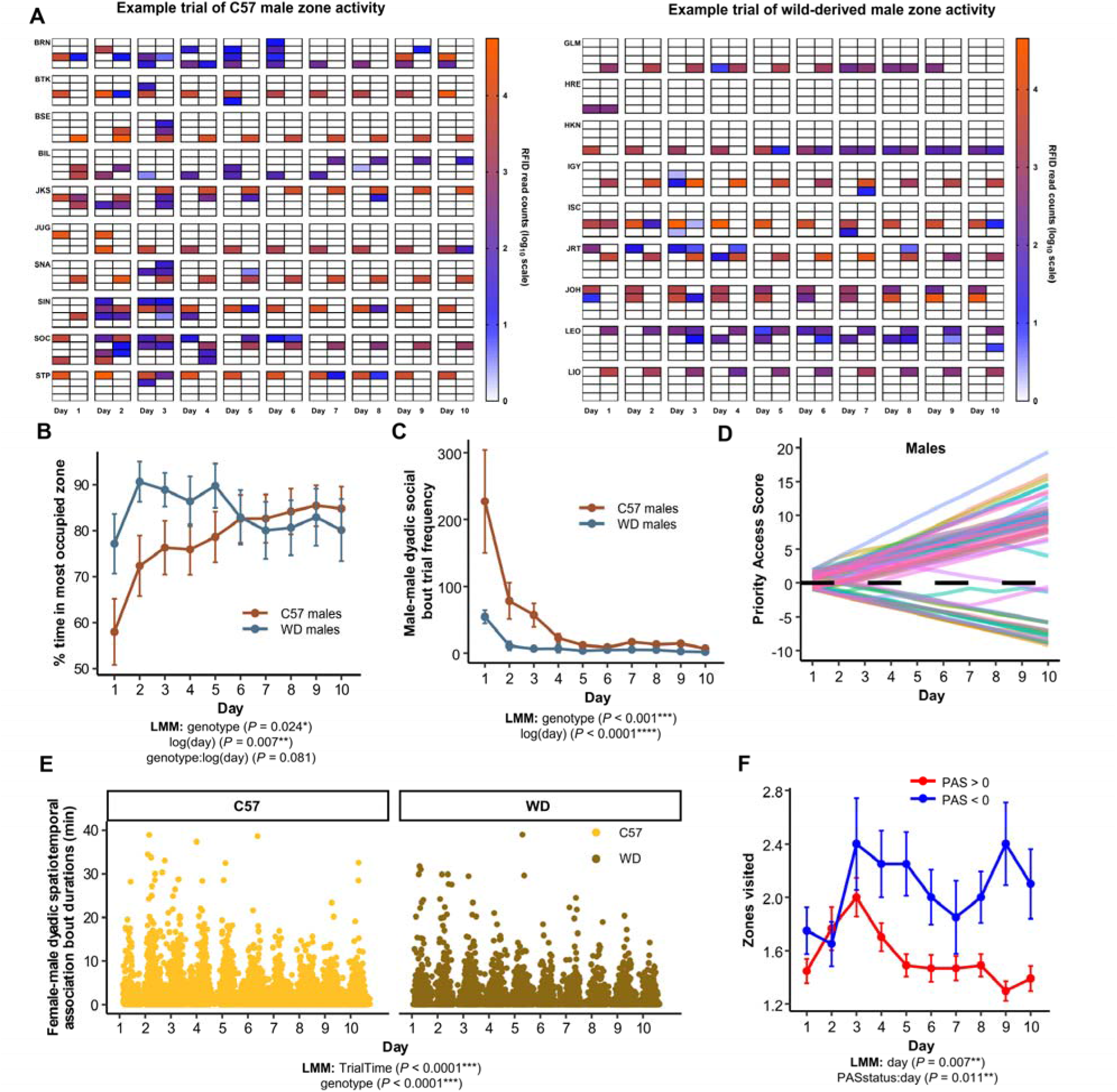
Territory establishment in C57 and wild-derived male mice. **(A)** Schematic of the resource zone locations (colored boxes) within the field enclosures (2×4 grids) showing patterns of male resource zone usage (rows) for an example C57 (left) and WD (right) trial across 10 days of activity (columns). White boxes indicate resource zones that were not visited by the focal individual. **(B)** Percentage of the total time a mouse was observed across all zones spent in a mouse’s top occupied zone (resource zones rank ordered by mouse occupancy time). **(C)** Male-male social grouping events fell rapidly over time and were less frequent in WD trials compared to C57 trials. **(D)** Over time males tend to either gain priority access to resource zones (i.e., are territorial) or do not (non-territorial males). Y-axis shows the evolving daily Priority Access Scores over 10 days of observation for males of both genotypes (See Methods for additional details on calculation of the daily PAS value). **(E)** C57 mice had longer male-female social grouping bout durations overall. For visualization purposes, the y-axis is cut off at 40 (n = 15,891 events shown out of 15,898 total events). **(F)** Daily zones visited were lower for high status males (Day 10 PAS > 0) than for low status males (Day 10 PAS < 0). Data are plotted as means ± s.e.m.

**Figure S3:**
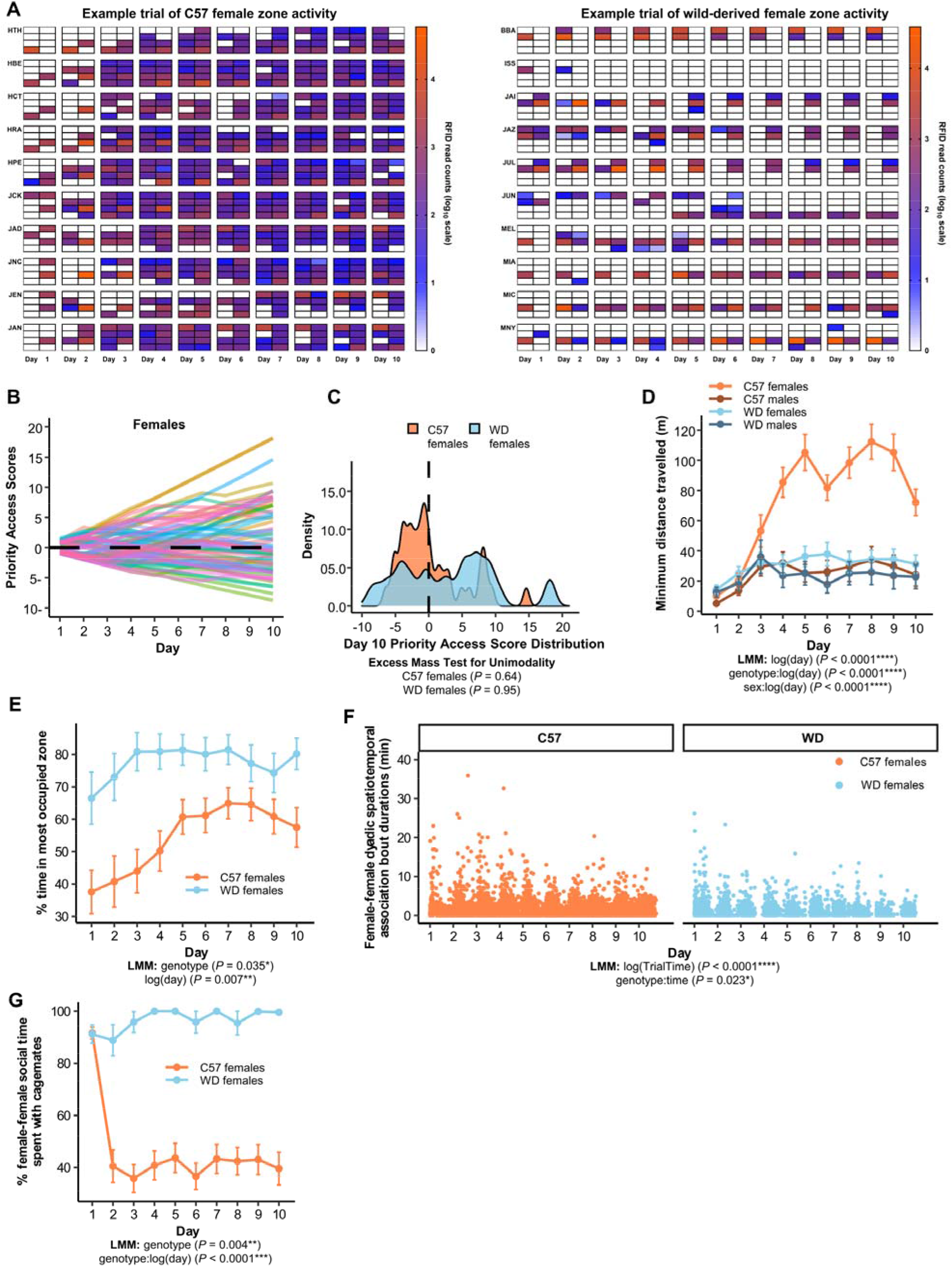
Distinct patterns of space utilization in C57 females. **(A)** C57 females extensively explore the available resource zones over the course of the trial compared to WD females. Example full trial data of female zone use in C57 (left) and WD (right) mice. Schematic of the resource zone locations (colored boxes) within the field enclosures (2×4 grids) showing patterns of zone usage for animals (rows) across 10 days of activity (columns). White boxes indicate resource zones that were not visited by the focal individual. **(B)** Daily Priority Access Scores over 10 days of observation for female mice. **(C)** Distributions of Day 10 Priority Access Scores for female mice are not multi-modal (excess mass test for unimodality from the multimode package), indicating decreased or inconsistent monopolization of resource zones amongst females. Higher scores indicate the extent to which a mouse maintained majority access over one or more resource zones relative to same-sex conspecific competitors (see Methods for details). **(D)** C57 female mice (n=40) differed from C57 male (n=40) and WD male (n=29) and female (n=30) mice in their estimated minimum distance travelled over the course of 10 days. **(E)** WD females spent more time in their most occupied zone than C57 females. **(F)** Female-female social grouping bout durations over time. For visualization purposes, the y-axis is cut off at 40 (n = 11,928 events shown out of 11,933 total events). **(G)** WD females spent nearly all of their female-female social time with cage mates after day 2, in contrast with C57 females who generally spent less than half of their female-female social time with cage mates. Data are plotted as means ± s.e.m.

**Figure S4:**
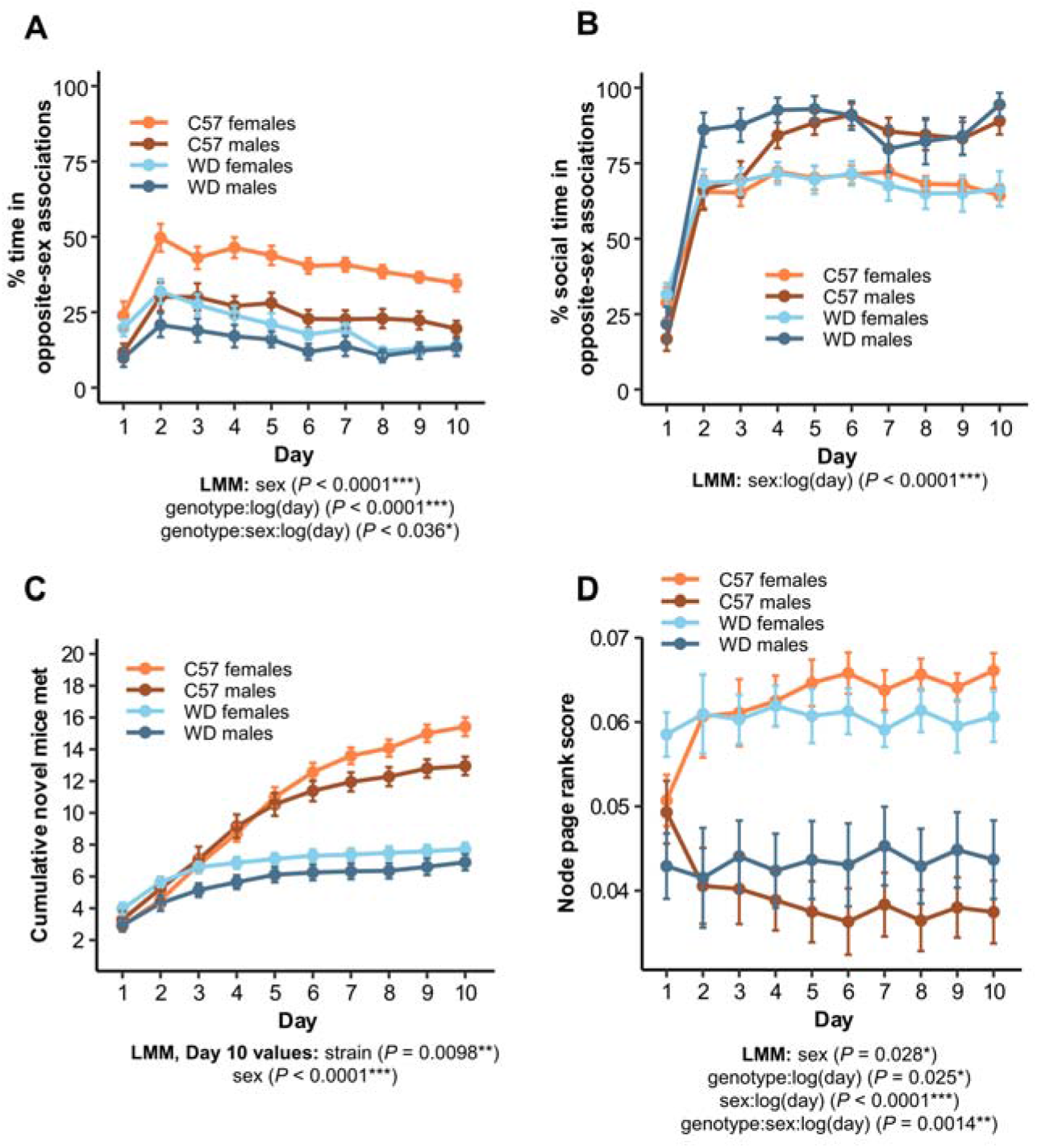
Repeatable differences in C57 social structure and network level properties over time. **(A)** Percentage of total recorded observation time spent in opposite-sex associations is higher in C57 mice over the course of the trial. **(B)** Percentage association time spent in opposite sex associations is higher in males than females for both genotypes. **(C)** C57 mice met a majority of the available novel social partners by the final day of the trial, while outbred mice did not. **(D)** Both C57 and outbred females exhibited high page rank scores relative to males of either genotype, indicating that females serve as major social connections through the network in both genotypes. Data are plotted as means ± s.e.m.

